# α-Synuclein-induced Kv4 channelopathy in mouse vagal motoneurons causes non-motor parkinsonian symptoms

**DOI:** 10.1101/856070

**Authors:** Wei-Hua Chiu, Lora Kovacheva, Ruth E. Musgrove, Hadar Arien-Zakay, James B. Koprich, Jonathan M. Brotchie, Rami Yaka, Danny Ben-Zvi, Menachem Hanani, Jochen Roeper, Joshua A. Goldberg

## Abstract

No disease modifying therapy is currently available for Parkinson’s disease (PD), the second most common neurodegenerative disease. The long non-motor prodromal phase of PD is a window of opportunity for early detection and intervention. However, we lack the pathophysiological understanding to develop selective biomarkers and interventions. By developing a mutant α-synuclein selective-overexpression mouse model of prodromal PD, we identified a cell-autonomous selective Kv4 channelopathy in dorsal motor nucleus of the vagus (DMV) neurons. This functional remodeling of intact DMV neurons leads to impaired pacemaker function *in vitro* and *in vivo*, which in turn reduces gastrointestinal motility which is a common, very early symptom of prodromal PD. We show for the first time a causal chain of events from α-synuclein via a biophysical dysfunction of specific neuronal populations to a clinically relevant prodromal symptom. These findings can facilitate the rational design of clinical biomarkers to identify people at risk for PD.

## Introduction

α-Synuclein is an established causal driver of PD: multiplications or point mutations in the *SNCA* gene lead to familial forms of PD^1,2^; single nucleotide polymorphisms near its locus increase the risk for idiopathic PD^3^; and α-synuclein aggregates are a main constituent of the Lewy pathologies (LPs) that are a hallmark of PD and other synucleinopathies^4^. These discoveries have driven extensive research into developing therapies aimed at either silencing the *SNCA* gene, reducing α-synuclein production, preventing α-synuclein aggregation, promoting degradation of intracellular and extracellular α-synuclein or preventing its propagation to other cells^5,6^. However, the lessons from Alzheimer’s disease clinical trials targeting betα-amyloids with similar strategies^7^ indicated that early detection before manifestation of clinical core symptom might also be essential for PD^5,6^. Therefore, for future disease-modifying therapies to be successful they will require early detection of the α-synucleinopathies, which in most (but not all) cases appear in the brainstem years before clinical diagnosis^8–11^. During this prodromal period, future PD patients already begin to suffer from a variety of non-motor symptoms (NMS), which can precede clinical diagnosis by decades^11^. It stands to reason that particular NMS are associated with the appearance of LPs in particular anatomical regions. For example, constipation and dysautonomia^12–14^ are likely related to the appearance (in Braak stage I)^15^ of misfolded α-synuclein in the medullary dorsal motor nucleus of the vagus (DMV) that innervates the gastrointestinal (GI) tract^16^. However, it is unknown whether dysautonomia is caused simply by DMV cell loss, or driven by LP-induced functional changes in surviving DMV neurons. This difference has important implications, as PD-selective DMV pathophysiology is likely to facilitate early detection compared to more generic phenotypes caused by cell loss and neuroinflammation. In the present study, we test this hypothesis by generating an adult onset mouse model of a selective medullary α-synucleinopathy. We use this model to identify an α-synuclein-induced channelopathy in DMV neurons that causes prodromal dysautonomia.

## Results

### Selective induction of α-synucleinopathy in vagal motoneurons of adult mice reduces vagal output and gastrointestinal motility

To determine whether α-synucleinopathy restricted to the medulla reproduces dysautonomia characteristic of prodromal PD, we injected adeno-associated viruses (AAVs) harboring either the mutated human *A53T-SNCA* gene (AAV-A53T) or an empty vector (AAV-EV) into the right cervical vagus nerve of C57BL/6JRccHsd mice (**Fig. 1a**). Six weeks later, the A53T α-synuclein transgene product was observed bilaterally within afferent sensory fibers of the medullary nucleus of the solitary tract (NST) and area postrema^17^ (**Fig. 1b**) as well as within ipsilateral cholinergic motoneurons of the DMV (**Fig. 1b,c**). Unbiased stereological analysis at this timepoint demonstrated that AAV injections *per se* – whether α-synuclein-expressing or empty vectors - induced mild cell loss with a median of approximately 20% – presumably due to mechanical damage to the vagus nerve fibers (**Supplementary Fig. 1**). Thus, DMV cell loss alone cannot explain divergent phenotypes in this mouse model.

**Figure 1.**
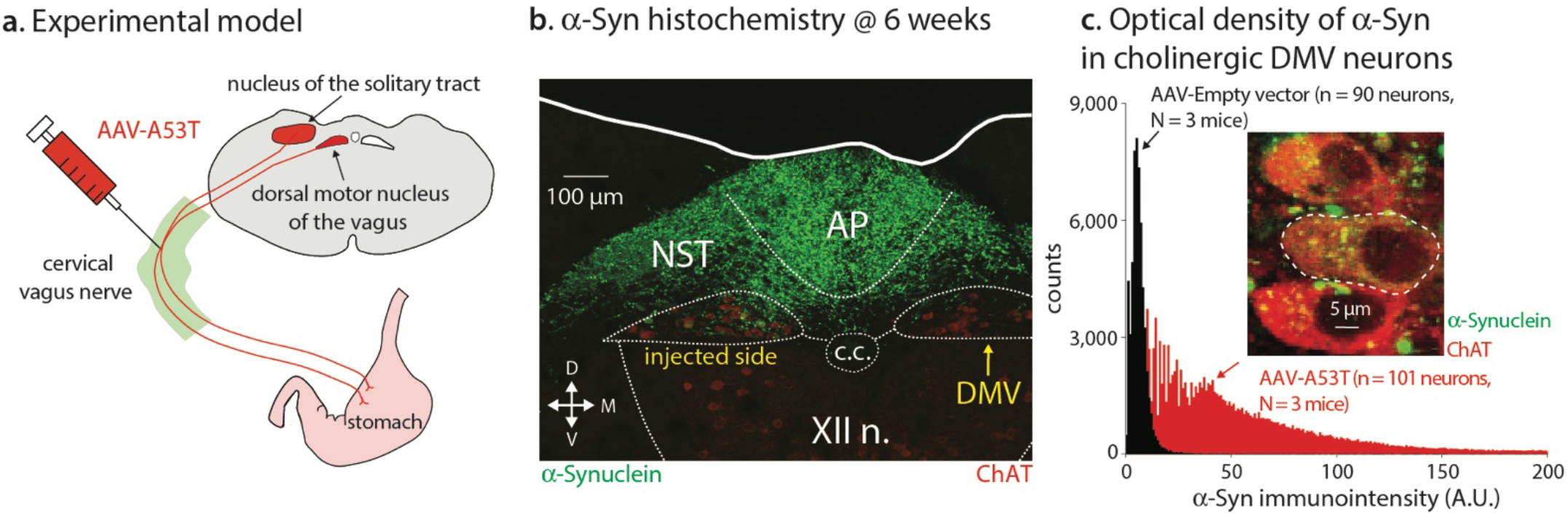
Model of adult onset α-synucleinopathy in the mouse dorsal medulla. (**a**) Injection of adeno-associated viruses (AAVs) harboring the mutated human *A53T-SNCA* gene into the cervical vagus causes the transfection of the dorsal motor nucleus of the vagus (DMV) and other regions such as the nucleus of the solitary tract (NST). (b) Six weeks post injection, α-synuclein (green) is visible in the area postrema (AP), in the NST and in the DMV, where immunohistochemical staining for choline acetyltransferase (ChAT) can be seen (red). XII n. – hypoglossal nucleus; c.c. – central canal. (**c**) Optical density measurements of the intensity of human α-synuclein fluorescent immunolabeling (green in inset) within regions-of-interest (dotted white line in inset) marked around the somata of ChAT-expressing neurons (red in inset) reveals a rightward shift in the distribution of immunointensity values from AAV-A53T-transfected relative to AAV-Empty Vector (AAV-EV)-transfected vagal motoneurons.

Six weeks after bilateral injection of the cervical vagus nerves, the AAV-A53T-injected mice produced larger stool than the AAV-EV injected mice (**Supplementary Fig. 2**), which is suggestive of slowed GI motility. Direct measurements of time of passage confirmed a slowing of GI motility at 6, but not 3, weeks after transfection (**Fig. 2a**). Using excised segments of the ileum in an organ bath, we ruled out that the slowed motility was due to a direct intestinal action of α-synuclein, despite its expression in vagal terminals (data not shown)^18^. We found that neither the responses of the ileal muscle to the muscarinic cholinoreceptor agonist pilocarpine, nor the frequency and amplitude of spontaneous contractions were altered by α-synuclein (**Fig. 3**). Similarly, responses of the ileal muscle to electric field stimulation (EFS) of the nerves were not altered (data not shown). Thus, because GI tract smooth muscles, pacemakers and nerves were intact in AAV-A53T treated mice, we hypothesized that the expression of mutant α-synuclein in the DMV reduced GI motility by inhibiting the electrical activity of DMV motoneurons, that normally maintains proximal gastric tone which augments GI motility^19,20^. Indeed, selective, unilateral transfection of cholinergic DMV neurons with AAVs harboring the inhibitory Designer Receptor Exclusively Activated by Designer Drugs (DREADDs), hM4Di, chemogenetically inhibited their electrical activity (**Supplementary Fig. 3)** – and was sufficient to recapitulate the slowed motility induced by AAV-A53T when the mice were intraperitoneally injected with the DREADD agonist clozapine-*N*-oxide (CNO), but not with vehicle. Moreover, injection of CNO to mice that were not injected with DREADDs had no effect on GI motility (**Fig. 2b**).

**Figure 2.**
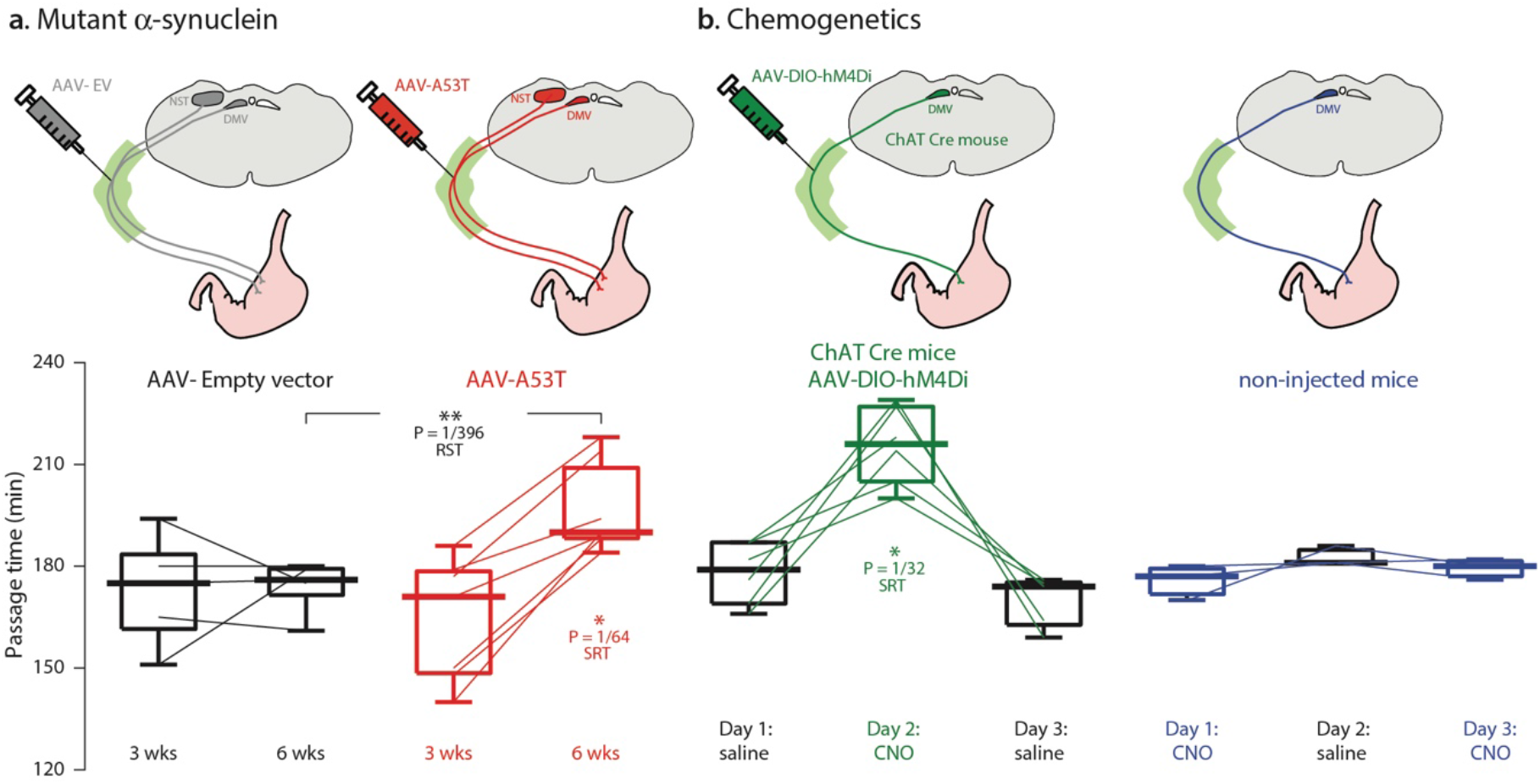
Gastrointestinal motility is slowed in the mouse model of medullary α-synucleinopathy. (**a**) Animals transfected with AAV-A53T but not with AAV-EV exhibited slowed GI motility at 6, but not 3, weeks after transfection. (**b**) Injection of AAVs harboring Cre-dependent hM4Di DREADDs into the cervical vagus of ChAT-Cre mice leads to selective transfection of cholinergic motoneurons in the DMV. Intraperitoneal injection of the DREADD ligand, clozapine-*N*-oxide (CNO), but not vehicle (saline) slows motility, indicating that silencing these neurons suffices to recapitulate the effect of AAV-A53T transfection. Injection of CNO had no effect on GI motility in non-injected mice. RST – two-tailed Wilcoxon Rank-Sum test; SRT – two tailed Wilcoxon Signed-Rank test.

**Figure 3.**
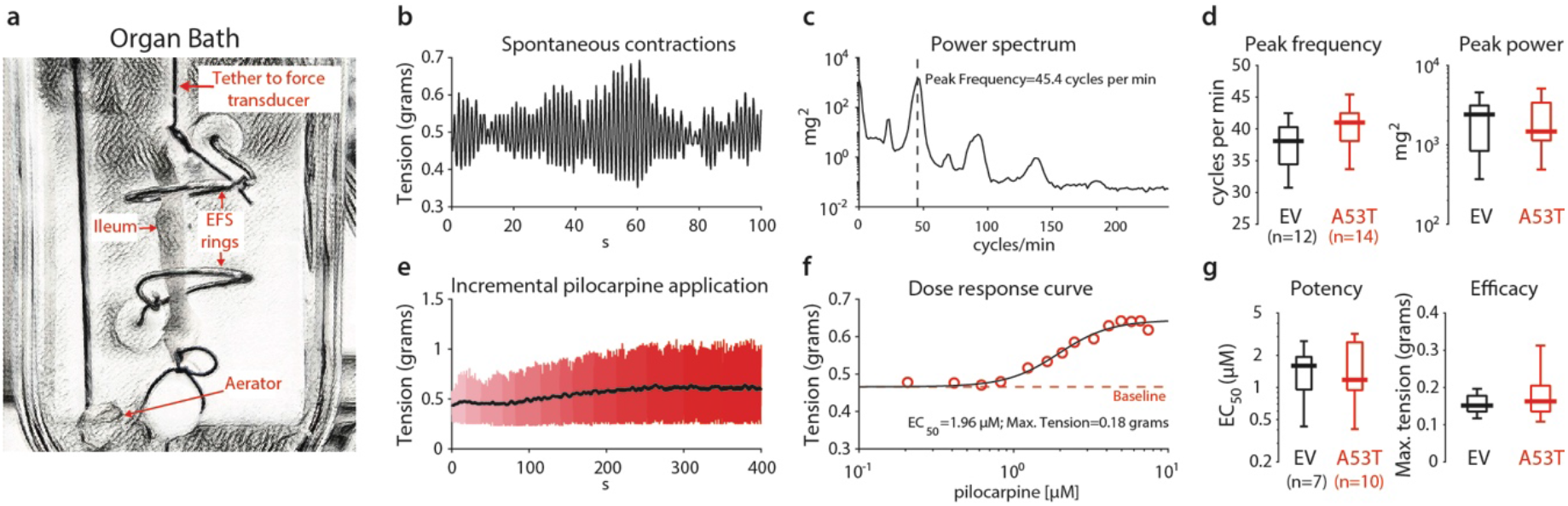
Intrinsic motility and susceptibility to muscarinic stimulation of excised ileal segments in an organ bath after induction of adult onset medullary α-synucleinopathy. (**a**) Organ bath used for the excised ileum preparation. Short segment of ileum is immersed into warm (37°C), oxygenated Krebs solution, while tethered to a force transducer, and surrounded by two ring electrodes for electric field stimulation (EFS). (**b**) Tension as a function of time, reveals spontaneous polymorphic oscillations in the excised segment. (**c**) Power spectrum of the spontaneous contractions reveals multiple spectral peaks. (**d**) Peak frequency of oscillations (extracted from the fundamental mode of the power spectrum) and amplitude of oscillations in the mice transfected with AAV-A53T (N=3 mice) or AAV-EV (N=2 mice) is similar. (**e**) Tension as a function of time during incremental increases in bath concentration of the muscarinic agonist, pilocarpine. (**f**) Dose response curve of pilocarpine extracted from panel E. (**g**) The potency (EC50) or efficacy of the direct effect of pilocarpine on ileal muscle in the mice transfected with AAV-A53T (N=2 mice) or AAV-EV (N=2 mice) is similar. Lowercase n’s indicate number of ileal sections used.

To determine whether vagal motoneuron firing rates were indeed slower in the AAV-A53T injected mice, we performed *in vivo* single-unit extracellular recording of DMV neurons under isoflurane anesthesia^21,22^. Using juxtacellular labeling combined with immunohistochemistry (**Fig. 4a**), 65% (17/26) of the recorded cells were positively identified as cholinergic DMV neurons (**Fig. 4b**), with the remainder being identified within the DMV via electrolytic lesions. The median firing rate of vagal motoneurons transfected with AAV-A53T was 60% slower than those transfected with AAV-EV. This reduction in firing rate was selective to vagal motoneurons, as neurons in the neighboring NST, where mutant α-synuclein was also expressed (**Fig. 1b**), discharged at control rates (**Fig. 4b**).

**Figure 4.**
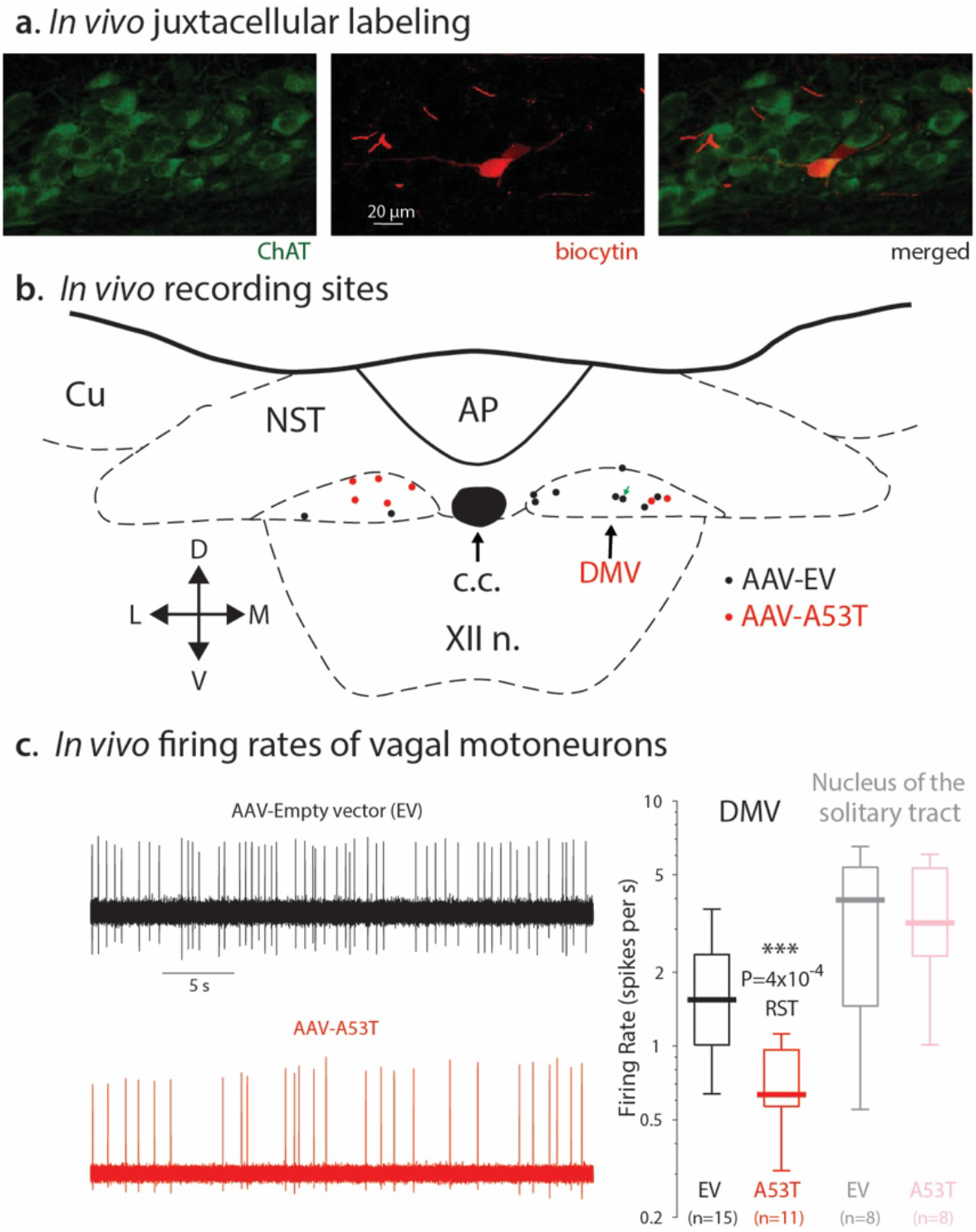
*In vivo* extracellular recording and juxtacellular labeling of cholinergic DMV neurons in the mouse model of medullary α-synucleinopathy. (**a**) Left: a ChAT-expressing (green) DMV neurons. Middle: juxtacellularly labeled DMV neuron red) after *in vivo* extracellular recording from an anesthetized mouse injected with AAV-A53T. Right: Merge of both images. (**b**) Map of recording sites of juxtacellularly labeled neurons (black and red dots correspond to AAV-EV and AAV-A53T transfected neurons, respectively). Green arrow indicates location of neuron from panel A. Abbreviations: AP – area postrema; c.c. – central canal; Cu – cuneate nucleus; DMV – dorsal motor nucleus of the vagus; NST – nucleus of the solitary tract; XII n. – hypoglossal nucleus. (**c**) Left: examples of extracellular recordings from mice injected with an AAV-EV (black) or with AAV-A53T (red). Right: box plot of firing rates demonstrates that AAV-A53T transfection leads to a reduction in the firing rates of DMV (N=5 EV- and N=8 A53T-injected mice), but not NST, neurons (N=5 EV- and N=4 A53T-injected mice). RST – two-tailed Wilcoxon Rank-Sum test.

### α-Synucleinopathy in mouse vagal motoneurons slows their autonomous pacemaker frequency due to elevated surface density of functional Kv4 channels

DMV motoneurons are autonomous pacemakers^23^, meaning that they discharge independently of synaptic input due to their intrinsic pacemaker currents. Thus, the most parsimonious explanation for the reduced *in vivo* firing rates in the AAV-A53T injected mice is that their intrinsic pacemaker frequency is lowered. To test this, we first measured in the whole-cell configuration the frequency-current relationship (f-I curve) of DMV motoneurons from acute medullary slices 6 weeks after AAV injection. The f-I curve characterizes neuronal excitability across the entire dynamic range of the neuron’s input-output relationship. We found that the f-I curve of DMV motoneurons from AAV-A53T injected mice was shifted to the right indicating a reduction in excitability in comparison to controls (**Fig. 5a**). In particular, the measurement in the zero-current condition demonstrated that the autonomous firing rate of DMV motoneurons from the AAV-A53T injected mice was reduced. In order to quantitatively compare the reduction in the autonomous *in vitro* firing rate with that observed *in vivo*, we repeated the measurement of the autonomous firing rate in the extracellular cell-attached configuration (that maintains the neurons’ metabolic integrity), and found that the autonomous firing rates were reduced in AAV-A53T-injected mice by 60%, relative to controls (**Fig. 5b**). The close agreement with the *in vivo* recordings (**Fig. 4a**) suggests that slowing of vagal motoneuron discharge in the intact animal is due mainly to a cell-autonomous change in excitability (while not ruling out additional circuit adaptations).These findings demonstrate that the ability of either intrinsic or extrinsic currents to depolarize the membrane potential of DMV motoneurons is curtailed by some α-synuclein-induced modification to their membrane biophysics (i.e., to an ionic current that underlies their slow autonomous pacemaking).

**Figure 5.**
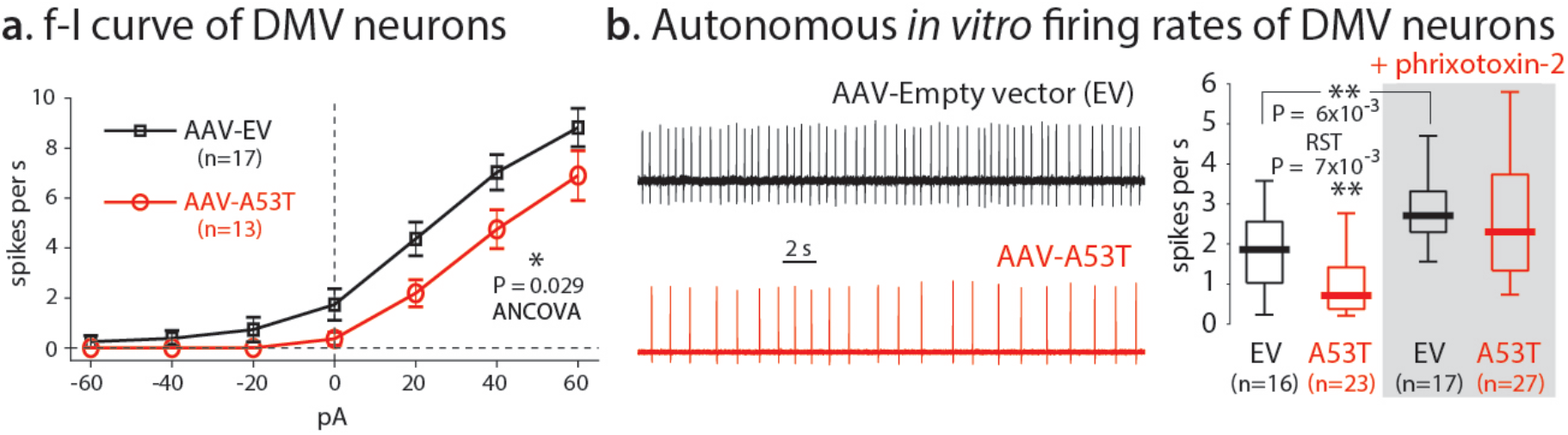
Reduction of pacemaker frequency and intrinsic excitability of DMV neurons in the mouse model of medullary α-synucleinopathy is caused by a dysregulation of Kv4 currents. (**a**) Frequency-intensity (f-I) curves of DMV neurons transfected with AAV-A53T (N=5 mice) are right-shifted relative to control (N=7 mice) indicating reduced excitability. (**b**) Examples of cell-attached recordings of DMV neurons that were transfected with either AAV-EV (black) or AAV-A53T (red) in acute brain slices. Traces are the time-derivative of passive current clamp recordings. Right: box plot of firing rates demonstrates a slowing of the pacemaker frequency in DMV neurons transfected with AAV-A53T (N=5 EV- and N=9 A53T-injected mice) that is equivalent to the reduction observed *in vivo* and that this reduction is prevented by preincubation of the slices in 1.3 μM phrixotoxin-2, a Kv4 selective antagonist (N=2 EV- and N=3 A53T-injected mice). RST – two-tailed Wilcoxon Rank-Sum test.

One ionic current that curtails the rate of membrane depolarization and the firing rate of DMV neurons is the α-type, fast inactivating voltage dependent K^+^ current carried by Kv4 channels^24,25^. Functional Kv4 channels act like a brake on spontaneous subthreshold depolarizations and slow down pacemaker frequencies. To test whether Kv4 channels were responsible for the α-synuclein-induced reduction in firing rates, we measured the autonomous firing rate of DMV neurons in the presence of 1.3 μM phrixotoxin-2, a Kv4 selective neurotoxin^24^. Phrixotoxin-2 prevented the reduction in firing rates in the AAV-A53T-injected mice relative to controls (and elevated the basal rate under both conditions, as reported previously)^24^ thereby proving that the firing rate reduction was entirely due a dysregulation of the Kv4 current.

To investigate the source of the Kv4 current dysregulation, we compared the Kv4 current biophysics in DMV motoneurons treated with either AAV-A53T or AAV-EV (**Fig. 6a**). While the voltage dependencies and kinetics of the A-type current in DMV neurons did not differ between the AAV treatments (**Fig. 6b**), the surface density of the Kv4 current was significantly higher in mice injected with AAV-A53T (**Fig. 6c**). The constitutive Kv4 window current in the subthreshold range (**Fig. 6b**), that is encountered during the spike afterhyperpolarization, prolongs the membrane depolarization and the lingering in the subthreshold range^24^ which suffices to explain the reduced pacemaker frequency. Importantly, our data argue against an upregulation of functional Kv4 channels, because the total whole-cell Kv4 current did not differ between the AAV treatments (data not shown), pointing to an alternative mechanism.

**Figure 6.**
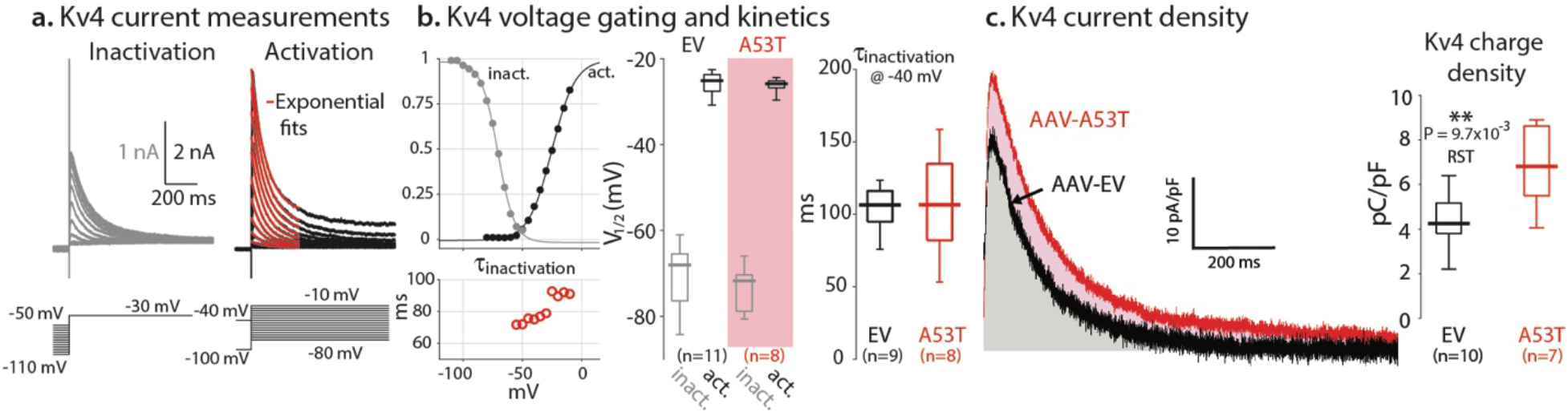
Surface density of Kv4 currents is elevated without changes to their voltage gating and kinetics. (**a**) Voltage clamp measurements of A-type currents yield voltage activation and inactivation curves and time constants of inactivation. Note that a window current flows constitutively in the subthreshold voltage range where the voltage activation and inactivation curves overlap. (**b**) Comparison of voltage dependency of activation and inactivation and channel inactivation kinetics between AAV-EV (N=6 mice) and AAV-A53T (N=4 mice) transfected mice reveals no difference. (**c**) Left: example of measurements of the inactivating A-type current density (depolarization to −40 mV), that is carried by Kv4 channels, from DMV neurons transfected with either AAV-EV (black, N=6 mice) or AAV-A53T (red, N=4 mice). Areas under the curves in gray and pink, respectively, indicate the Kv4 charge densities, which are significantly higher in the DMV neurons transfected with AAV-A53T (box plots). RST – two-tailed Wilcoxon Rank-Sum test.

### An α-synuclein-induced, cell-autonomous shrinkage of vagal motoneurons causes a selective dysregulation of Kv4 current densities

α-Synuclein overexpression has been shown to cause vagal motoneurons^26^ and other neuronal types^27^ to shrink. Indeed, measurement of the long axis of the somata of DMV neurons from animals transfected with AAV-A53T demonstrated that it was shorter in human α-synuclein-positive DMV neurons in comparison to neighboring human α-synuclein-negative DMV neurons (**Fig. 7a**). Note that human α-synuclein-negative neurons in AAV-A53T transfected mice were not different in size compared to controls in the AAV-EV transfected mice (**Fig. 7a**). This indicates that shrinkage of cell size was due to a cell-autonomous process, and gave – at least at the timepoint of analysis - no evidence for the contribution of additional non-cell-autonomous conditions, such as neuroinflammation^26,28^. To better gauge the degree of reduction in surface area, we filled DMV neurons from both AAV-EV and AAV-A53T transfected mice with biocytin. We then reconstructed their perisomatic region using fluorescent confocal microscopy to estimate their surface area. Using these morphological measurements, we found that the surface area was reduced by 38% from a median area of approximately 3,600 μm^2^ in the AAV-EV transfected mice to approximately 2,200 μm^2^ in the AAV-A53T transfected mice (**Fig. 7b**, left). These measurements were corroborated by an electrophysiological measurement of the membrane surface area. The whole-cell capacitance of a neuron scales with the somatic surface area because it is the product of the surface area and the specific capacitance of the bilipid membrane (C_m_)^29^. Measurements of the whole-cell membrane capacitance of the DMV neurons (**Fig. 7b**, right) demonstrated that it was reduced by 33% from approximately 32 pF in neurons transfected with AAV-EV to approximately 22 pF in DMV neurons transfected with AAV-A53T. Thus, there is a quantitative agreement between the A53T-SNCA induced reductions of surface area estimated by either electrophysiological or morphological methods. Moreover, in both AAV treatments, the ratio of the median whole-cell capacitance to the median surface area very closely reproduced the known value of C_m_ (1 μF/cm^2^)^29^. In summary, the Kv4 channelopathy in human α-synuclein-positive DMV neurons is likely caused by a mismatch of their Kv4 surface expression in shrinking DMV neurons.

**Figure 7.**
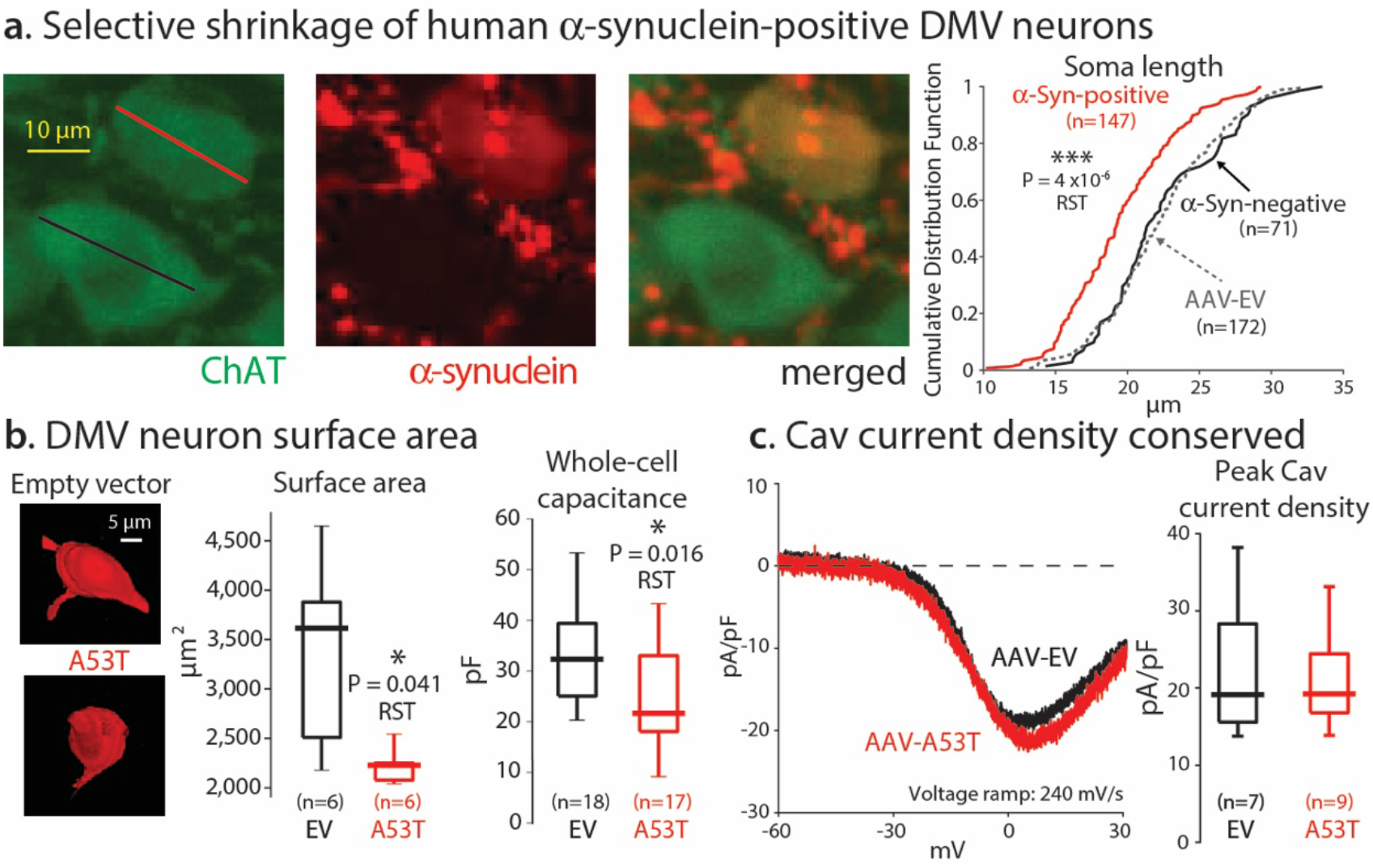
Selective shrinkage of human α-synuclein-positive DMV neurons. (**a**) Left panels: examples of the measurement of the long axis of the soma of DMV neurons from mice injected with AAV-A53T. Staining for ChAT and human α-synuclein, demonstrate the selective shrinkage of the human α-synuclein-positive (long axis marked in red), but not the neighboring human α-synuclein-negative neuron (long axis marked in black). Right panel: cumulative distribution function (cdf) of the lengths of human α-synuclein positive and negative neurons (N=3 A53T-injected mice). Dashed line indicates the cdf of neurons measured from the AAV-EV injected mice (N=2). (**b**) Left: Examples of biocytin filled DMV neurons from AAV-A53T and AAV-EV injected mice that were reacted with streptavidin conjugated to Alexa 594. Middle: box plots of the somatic surface areas of the biocytin-filled neurons as estimated with the FIJI software (N=5 EV- and N=4 A53T-injected mice). Right: whole-cell capacitance measurement of DMV neurons from mice transfected with either AAV-EV (N=10 mice) or AAV-A53T (N=11 mice). The morphological and physiological measurements demonstrate that only DMV neurons that express A53T α-synuclein shrink. (**c**) Left: examples of the Co^2+^-sensitive voltage-activated Ca^2+^ (Cav) current in response to fast voltage ramps from DMV neurons transfected with either AAV-EV (black, N=5 mice) or AAV-A53T (red, N=6 mice). Right: box plots of peak Cav current density, estimated by fitting activation curves to individual Co^2+^-sensitive current density measurements^30^ indicate that they are equivalent under both AAV treatments despite the shrinkage, due to a downregulation of the total Cav current in DMV neurons transfected with AAV-A53T (data not shown). RST – two-tailed Wilcoxon Rank-Sum test.

Because Kv4 channel expression did not adapt to the A53T-driven reductions in surface area, we asked whether this maladaptation was selective to Kv4 channels. To address this, we measured voltage activated Ca^2+^ (Cav) currents. We found that the total Cav currents in DMV neurons were reduced (data not shown), as shown previously in transgenic mice overexpressing human A53T α-synuclein^30^. Intriguingly, the reduction in Cav currents of DMV neurons in the AAV-A53T-transfected mice matched the reduction in their surface area. Thus, in contrast to Kv4 channels, Cav channels may exhibit intact homeostatic control (**Fig. 7b**). Similarly, measures of other electrophysiological properties of the DMV pacemaking (*e.g*., action potential threshold, width and afterhyperpolarization) were largely unaffected (**Supplementary Fig. 4**), ruling out significant changes to other voltage-activated channels involved in pacemaking^23,24,30,31^. Taken together, these findings suggest that the A53T-induced Kv4 channelopathy in DMV neurons is both selective and maladaptive.

In summary, we have shown that acute expression of mutated α-synuclein in DMV motoneurons of adult mice leads to a selective elevation in the surface density of Kv4 channels in a cell-autonomous fashion that is secondary to somatic shrinkage without cell loss. The elevated Kv4 density slows the depolarization to action potential threshold and thereby reduces the DMV motoneurons’ pacemaker frequency. The ensuing reduction in vagal parasympathetic tone results in slowed GI motility (**Figure 8**). This model provides a feasible pathophysiological mechanism for constipation in PD – one of its most common, early-onset prodromal symptoms. Elucidating the physiological underpinning of prodromal symptoms paves the way for a rational design of clinical biomarkers for early diagnosis of the PD, and neuroprotective strategies to combat it.

**Figure 8.**
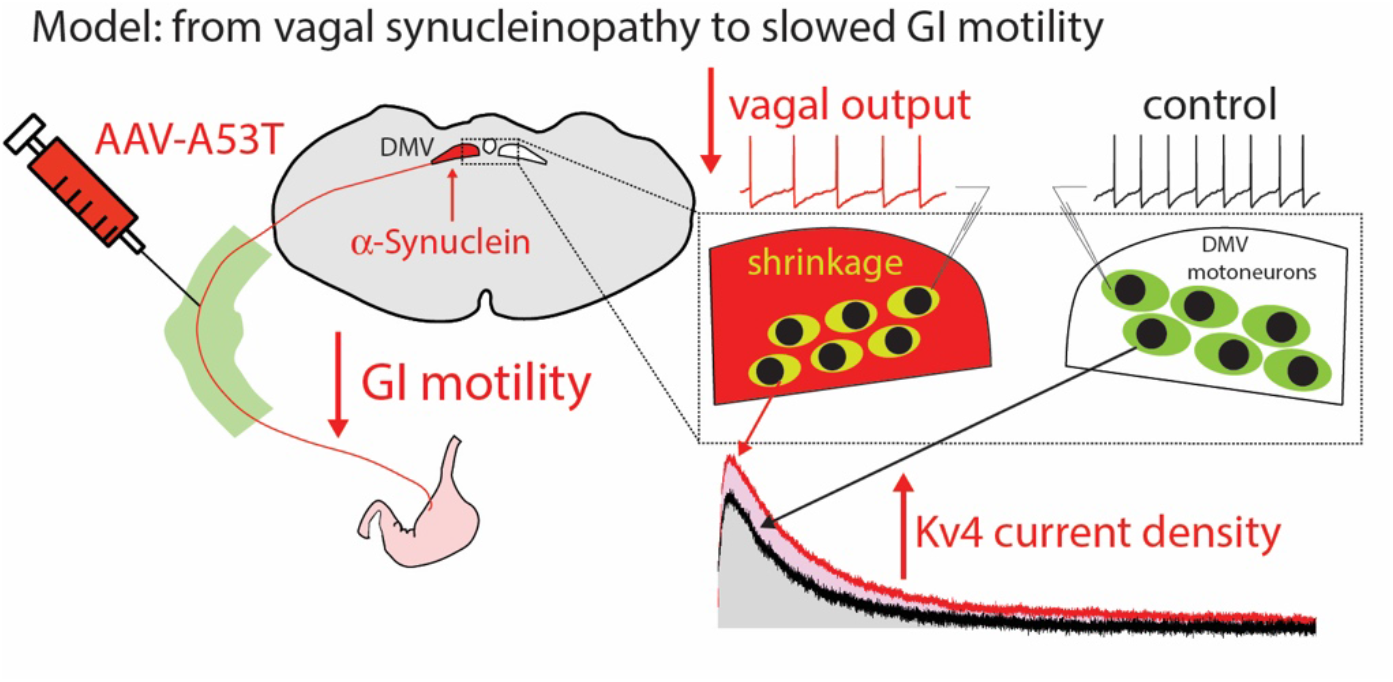
Working model of how the medullary α-synucleinopathy reduces GI motility. Expression of mutated α-synuclein in DMV motoneurons reduces vagal tone and GI motility due to a selective elevation in the surface density of Kv4 channels that is secondary to their shrinkage.

## Discussion

The notion that brain stem α-synucleinopathies are linked to prodromal NMS of PD, including dysautonomia (e.g., constipation) is not new^32–34^. Earlier studies have even proposed that clinical autonomic measures may be sensitive enough to gauge the degree of brain stem LPs^35^. However, it is unknown whether it is α-synuclein-induced cell loss in brain stem nuclei like the DMV^36,37^ that is responsible for the dysautonomia. Functional changes induced by α-synuclein might also contribute to vagal dysfunction and might even precede cell-loss mediated impairments. These functional changes can be divided into cell-autonomous effects (e.g. altered physiology of DMV neurons that harbor α-synuclein pathology)^38^ and non-autonomous effects (e.g. involvement of surrounding microglia leading to neuroinflammation)^39^. In contrast to the strong evidence for neuroinflammation and cell loss in prodromal PD^39^, we know nothing about functional changes in viable DMV neurons in humans and only very little in model systems^38^. In a global A53T-SNCA transgenic mouse model, we previously reported *in vitro* reductions in activity-dependent calcium loading and oxidative enhancement of Kv4 channels in DMV neurons^30^. In contrast, global transgenic expression of mutant α-synuclein led to *in vivo* and *in vitro* hyperexcitability with Kv4 dysfunction in dopamine substantia nigra neurons^40^. Importantly, these global transgenic models cannot capture prodromal PD states where α-synuclein pathology is restricted to a small number of sites. Based on immunohistochemical studies in human brains, Braak and colleagues have proposed a set of six stages of increasing expansion of LPs, which have been confirmed by others and may represent the most common spatiotemporal sequence of LPs^41,42^. Importantly, LPs in DMV are a hallmark of the earliest Braak Stage I^43^ and might be associated with various degrees of cell loss^36,37,39^. The demonstration of α-synuclein spreading throughout neural circuits in rodent models^41,42,44,45^ provided a plausible biological mechanism in support of the Braak hypothesis. Moreover, even processes that are currently believed to occur at the earliest stages, including the spread of α-synuclein from the periphery to DMV neurons were successfully modelled in rodents^41,42,46^. However, as potential functional implications are still unknown, we expanded these prodromal models to include functional physiological studies *in vitro* and *in vivo*.

In our model, 6 weeks after injection of AAV-A53T into the cervical vagus, the α-synuclein immunoreactivity was restricted to fibers, but not somata, in the NTS and area postrema. These fibers belong to afferent sensory fibers whose somata are located outside the cranium. Thus, the α-synuclein expressed in the afferent fibers seems not to cross trans-synaptically into their target neurons in the NST and area postrema. Similarly, α-synuclein taken up by efferent (including DMV motoneuron) fibers does not cross trans-synaptically into the GI tract^18^, leaving, as we demonstrated here, the intrinsic physiological properties of the GI nerves and muscles intact, as recently observed in another rodent model of PD^47^. Also, our stereological analysis demonstrated that α-synuclein expression in DMV neurons did not induce cell death *per se*. Thus, our model is well suited to capture a very early timepoint in prodromal PD, where the phenotype is mostly driven by the pathophysiology of DMV neurons. Indeed, we have demonstrated that a selective physiological dysregulation of Kv4 channel density in DMV neurons causes a reduction in vagal output, which slows GI motility and elevates HR.

We have shown previously that DMV neurons in transgenic mice that globally overexpresses human A53T α-synuclein undergo cell-autonomous adaptations (compared with control mice) in the form of a transcriptional downregulation of Cav currents, which leads to lower basal mitochondrial oxidative stress and improved oxidative function of Kv4 channels. The latter effect was only revealed through the dialysis of an antioxidant into the cell via the patch pipette, because no difference was found in the amplitude, voltage dependence or kinetics of the Kv4 currents^30^. The current study was therefore motivated by the concern that these observed adaptations in expression and oxidative function of Cav and Kv4 channels reflected some developmental adaptation in the transgenic mouse model, rather than an adaptive response that plausibly represents the disease process in adult PD patients. Therefore, in the current, adult onset brain stem-selective α-synucleinopathy model, we initially set out to replicate the Cav and Kv4 findings.

We were able to replicate the reduction in total Cav current, which suggests to us that mutant α-synuclein drives the downregulation of Cav channels robustly regardless of whether it accumulates acutely or developmentally. Because basal mitochondrial oxidative stress in DMV neurons requires Ca^2+^ influx via Cav2 channels, the Cav2 current downregulation reduces basal oxidative stress in DMV motoneurons^23,30^. Therefore, this response can be considered a protective response, and may explain why DMV neurons are relatively spared – according to some reports^36^ - compared to other neurons that are vulnerable in PD. Interestingly, when normalizing the total Cav current to a current of density, we found that in the adult onset α-synucleinopathy model, current density is preserved despite the shrinkage. While this might be a sheer coincidence, the normalization may represent a controlled homeostatic mechanism which titers the degree of channel downregulation to match the shrunken cell size. Either way, because high-voltage activated Cav2 currents measured in DMV neurons flow primarily as a consequence of spiking, they do not strongly affect pacemaking frequency^24^. Indeed, we have seen in the A53T transgenic mice that the pacemaking frequency is unaltered despite the Cav2 downregulation^30^.

In contrast to the Cav2 currents, the total Kv4 currents were unchanged in the adult onset α-synucleinopathy model leading to an effective upregulation of Kv4 current density, which was secondary to the cell shrinkage. Neither the shrinkage nor the Kv4 current density upregulation were observed in the A53T transgenic mouse. Moreover (again in contrast to the Cav2 currents) the pacemaker frequency of DMV neurons is quite sensitive to the amplitude of the Kv4 currents (**Fig. 5b**)^24^. Accordingly, in transgenic A53T mice where there is no change in the size of the Kv4 currents there is no change in pacemaker frequency^31^; whereas in the current model the upregulation of the Kv4 current density reduces the autonomous firing rate. While it would have been nice to provide independent proof that upregulating Kv4 channel density *per se* selectively in DMV motoneurons slows GI motility, this is not a manipulation that can be achieved easily in a controlled manner *in vivo*. Nevertheless, we believe we provided compelling evidence using chemogenetics and a selective Kv4 channel antagonist for a causal chain of events leading from α-synucleinopathy, via the selective Kv4 channelopathy to the impaired GI motility (e.g., constipation) and dysautonomia that are prevalent in prodromal PD.

Finally, the fact that the action potential threshold, width and afterhyperpolarization are largely unchanged in the current model suggests that the persistent sodium, and the large- and small-conductance calcium activated potassium currents, respectively^23,30^, are either hardly affected by the shrinkage (e.g., arise from compartments such as the axon that may not be affected by the shrinkage) or undergo some adaptive regulation like the Cav currents. In summary, while some currents are unaffected by the shrinkage, and while Cavs currents seem to normalize their surface density, it is clear that Kv4 currents selectively fail to normalize in response to the α-synuclein-mediated shrinkage of DMV neurons, and this failure triggers the chain of events that leads to slowed GI motility.

We made use of human mutated A53T α-synuclein, as it is an established cause for familial PD^2,38^. While corroboration of our finding in additional animals models of prodromal PD (e.g., that overexpress wildtype α-synuclein^42,46^, or models of gut-to-brain transmission of synucleinopathies^41^) is warranted, the aim of the present study was not to generate a strictly Braakian recapitulation of the progression of pathology (which has already been firmly established in animal models^41,42,45^). Indeed our findings are orthogonal to the question of the spread of α-synuclein. Instead, we aimed to produce a selective synucleinopathy localized to the DMV, thereby replicating an early timepoint in PD progression. Although it has been established the brain stem synucleinopathies are associated with prodromal symptoms^34^, ours is – to the best of our knowledge – the first study to provide a causal link from α-synuclein via ion channel dysregulation and reduced neuronal activity *in vivo* to a prodromal non-motor symptom of PD. This establishes an additional perspective – cell-autonomous neuronal pathophysiology – in the etiology of PD to complement well-established pathological and molecular approaches. It also provides a novel set of questions for follow-up studies to understand which molecular and cell-biological mechanisms induce α-synuclein-mediated shrinkage and channel dysregulation^26,27^.

### Clinical implications

In our model, the induction of an α-synucleinopathy in DMV motoneurons caused shrinkage and dysregulated surface densities of Kv4 channels, which in turn reduced firing rates. This reduction in vagal tone effectively slowed GI motility. If a similar mechanism is operative in prodromal PD patients with impaired GI motility, the translational challenge would be to establish a selective functional test for prodromal patients or others at risk for PD. As we have identified Kv4 channels in DMV neurons as molecular targets, a selective pharmacological challenge would be conceptionally most promising. While there is currently no selective Kv4 channel modulator in the clinic, the non-selective Kv channel blocker 4-aminopyridine^48^, used in multiple sclerosis^49^ and Lambert-Eaton myasthenic syndrome^50^, could potentially be useful, although off-target effects may be inevitable. For readout of these tests, monitoring changes in GI motility is, in principle, possible. Even in the absence of selective Kv4 pharmacology, the new level of mechanistic understanding provided by our study could form the basis for an improved differential diagnosis of prodromal dysautonomia in PD.

## Acknowledgments

This work was supported by grants from the German-Israeli Foundation for Scientific Research and Development (no. I-1294-418.13/2015) to J.R. and J.A.G, the Collaborative Research center 815 “Redox Signaling” program to J.R, and the European Research Council (no. 646880) to J.A.G. We thank Prof. Aron Troen for help with the stereological analyses, and Engs. Eugene Konyukhov and Anatoly Shapochnikov for excellent technical support.

## Author Contributions

Conceptualization, W-H.C., J.R. and J.A.G.; Methodology, W-H.C., R.E.M., H.A-Z., D.B-Z., M.H., J.R. and J.A.G.; Formal Analysis, W-H.C., R.E.M. and J.A.G.; Investigation, W-H.C., L.K., R.E.M, H.A-Z.,D.B-Z., J.A.G.; Resources, J.B.K., J.M.B, R.Y, M.H., J.R. and J.A.G.; Data Curation, W-H.C, R.E.M., J.R. and J.A.G; Writing – Original Draft, W-H.C., R.E.M., J.R. and J.A.G; Writing – Review & Editing, W-H.C., L.K., R.E.M, H.A-Z., J.B.K., J.M.B., D.B-Z., M.H., J.R. and J.A.G.; Visualization, W-H.C., R.E.M, J.R. and J.A.G; Supervision, J.R. and J.A.G.; Project Administration, J.R. and J.A.G.; Funding Acquisition, J.R. and J.A.G.

## Methods

### Ethical Statement

All experimental procedures on mice adhered to and received prior written approval from the Hebrew University Institutional Animal Care and Use Committee and from the German Regierungspräsidium Darmstadt.

### Animals

All experiments were conducted on male C57BL/6JRccHsd mice except for the chemogenetic experiments (see below) which were conducted one homozygous male ChAT-IRES-Cre transgenic mice (Jackson Laboratory, stock no. 006410).

### Cervical vagus injections

6-7 week old mice (>25 grams) were anesthetized with an *i.p*. mixture of ketamine (100 mg/kg) and medetomidine (83 μg/kg) and ointment was applied to prevent corneal dying. Mouse was held in place in a supine position with paper tape, and temperature was maintained at 37°C with a heating pad. Animals were hydrated with a bolus of injectable saline (5 ml/kg) mixed with an analgesic (5 mg/kg carpofen). The hairs around the neck area were removed with an electric razor and hair removing gel. Before cutting the skin, 70% alcohol and betadyne were used to sterilize at that area. The central line of the skin was vertically cut with surgical scissors from the sternum area to the lower jaw area (app. 2cm long), and the thoracic glands were gently teased apart. For unilateral injections, the right cervical vagus was exposed from beneath the carotid (after careful separation from fat and connective tissue) and held away from the pulsations with a paper pointer. The nerve was pierced with a glass pipette pulled to a very fine tip (3-4 times thinner than the vagus nerve trunk), and adeno-associated virus (AAV) particles were injected with a Nanoject 3 (Drummond scientific) at a speed of 1 nl/sec for 30 secs injection with 30 secs pause for a total 33 cycles (i.e., a 990 nl in total). After the injection, the pipette was left for an extra 1 min before slowly being retracted. At this point, the injection step terminated, and the paper pointer was gently removed so as not to damage the nerve. For bilateral injections the process was repeated on the left side. Both glands were placed back above the neck region, and the skin was sutured. At the end, the betadyne and alcohol sterilization was repeated. The animal was woken up with Anti-Sedan (0.42mg/kg *i.p*.) and kept warm until it is awake (app. in 30 mins). Animals were monitored and received carpofen for 3 additional days.

### Adeno-associated viruses (AAVs)

The C57BL/6JRccHsd mice were injected with 5 × 10^12^ GC/ml of and AAVs that either harbors a human mutated A53T form of the α-Synuclein gene (AAV1/2-CMV/CBA-human-A53T-alpha-synuclein-WPRE-BGH-polyA) or the harbors an empty vector (EV) (AAV1/2-CMV/CBA-empty vector-WPRE-BGH-polyA) (Genedetect, new Zealand). The ChAT-IRES-Cre mice were injected with an AAV that harbors a Gi Designer Receptor Exclusively Activated by Designer Drug (DREADD) gene (AAV9-hSyn-DIO-hM4Di-mCherry, 8.8 × 10^12^ GC/ml, ELSC Vector core), which unlike in the case of the C57BL/6 JRccHsd mice only expressed in the cholinergic cells of the DMV and not in the NST.

### *In vivo* GI motility assay

One week before AAV transfection, the C57BL/6JRccHsd mice (>25 g) were lightly sedated with isoflurane until the breathing pattern reached to 1 cycle/sec Chiu et al. 2020 (observed from the chest movement). Mice were gavaged, between 10 and 11 am, with 0.3 ml of 6% carmine-red (Sigma, Germany) mixed with 0.5% methyl cellulose (Sigma, Germany) using a 22G feeding tube. After 2.5 hours, the animals were transferred into a clean cage with fresh beddings, in order to monitor the latency at which the first reddish feces pellets were observed. The gavage feeding was repeated twice per animal (once per day) to determine consistent and stable results. Mice that were inactive or asleep (approximately 15%) were excluded. The actual measurements of passage time were conducted 3 and 6 weeks after transfection.

### Stool mass measurement

Mice injected with either AAV-EV or AAV-A53T was housed individually between 10 and 11 am in a clean cage with fresh beddings. Two and a half hours later the stools were collected and dehydrated, and the total stool mass was measured.

### Chemogenetics

ChAT-Cre transgenic mice that were transfected with the AAV harboring the DREADDs were tested 5-6 weeks post transfection. Each animal was subjected to 3 measurements on 3 different days. On the first day, the animals were gavage and *i.p*. injected with. vehicle injection of 0.9% saline (10 ml/kg). Two days later, in order to activate the Gi DREADDS overexpressed in the DMV the mice received an identical gavage feeding with an *i.p*. injection of 3 mg/kg water-soluble clozapine-*N*-oxide (CNO, HelloBio, UK) dissolved in injectable 0.9% saline. Following an additional 2 days, the vehicle gavage was repeated. Non-transfected animals were used as controls, and underwent the same measurements except that the first and last measurements were with CNO injections, while the middle one was saline.

### *Ex vivo* GI motility assay

Mice were perfused transcardially with ice-cold 0.9% phosphate buffered saline (for histological verification of DMV transfection, see below) and the entire GI tract from between the diaphragm and cecum was transferred into ice-cold Krebs solution carbogenated with 95% O_2_–5% CO_2_ and containing (in mM): 118 NaCl, 4.7 KCl, 14.4 NaCHO_3_, 5 CaCl_2_ 1.2 MgSO_4_, 1.2 NaH_2_PO_4_, 11.5 glucose. Sections of the ileum (approx. 1 cm long) were immersed in an organ bath (20 ml volume) filled with warm (37°C), carbogenated Krebs solution, and tied with a silk suture to a force transducer. The transducer signal was logged onto a PC computer with a National Instruments A/D board using custom made code in LabView (National Instruments). Sections were given 30 minutes to accommodate during which spontaneous contractions were recorded and then were subjected to increasing concentrations on pilocarpine (Sigma). Electric field stimulation (EFS) was conducted with a Grass S44 Stimulator using 0.5-ms-long, 55V pulses at 1, 2, 5, 10, 20 an 30 Hz for a duration of 15 s. The current was passed between two rings that surrounded (but did not touch) the ileal section inside the organ bath. We used spectral analysis to characterize spontaneous contractions and fit Hill equations to the dose response curves of tension as a function of pilocarpine concentration or EFS frequency.

### *In vivo* recordings and juxtacellular labeling

Mice were anesthetized in a non-rebreathing system with isoflurane (induction 2.5%, maintenance 0.8–1.4% in O_2_, 0.35 l/min) and were placed in a stereotaxic frame (David Kopf) and eye lubricant (Visidic, Bausch and Lomb, Berlin, Germany) was applied to prevent corneal drying. Lidocaine/prilocaine ointment (Emla^®^ cream, Astra Zeneca, Wedel, Germany) was used as a local analgesic at the incision site. Body temperature (33–36°C), heart rate (5–10 Hz), and respiration (1–2 Hz) were constantly monitored by the vital sign oximeter (Mouse OX, STARR Life Sciences, USA). Bregma was set to be 1 mm lower than lambda, so that the exposed brain stem will lie in a horizontal plane. All the neck muscles attached on the occipital bone were removed, a small craniotomy (± 0.5 mm from midline) was bored at the caudal position −4.0 to −4.3 mm from lambda. Micromanipulator (SM-6, Luigs and Neumann) was used to lower the electrodes (1-2 μm /sec) to the recording site [For DMV – AP: −4.1 to −4.3 mm (from lambda), ML: ± 0.1 to 0.3 mm (from lambda), DV: −2.8 to −3.0 mm (from the surface of cerebellum; For NTS – same AP and ML, DV: −2.3 to −2.7 mm].

Glass electrodes (10–20 MΩ; Harvard Apparatus) filled with 0.5 M NaCl, 10 mM HEPES, 1.5% neurobiotin (Vector Laboratories) were used for recording. The extracellular singleunit activity was recorded for 1-3 mins and the signals were acquired with EPC-10 A/D converter (PatchMaster software, Heka; sampling rate 12.5 kHz for spike train analyses and 20 kHz for AP waveform analyses). The extracellular signals were amplified 1,000x (ELC-03M, NPI Electronics), notch- and bandpass-filtered 0.3–5 kHz (single-pole, 6 dB/octave, DPA-2FS, NPI Electronics). The signals were displayed both on an analog oscilloscope and via an audio monitor.

To identify the anatomical location and neurochemical identity of the recorded neuron, following extracellular single-unit recordings, the neurons were labeled with neurobiotin using juxtacellular in vivo labeling technique^51^. Microionophoretic current was applied (1-7 nA positive current, 200 ms on/off pulse), via the recording electrode with continuous monitoring of the firing activity. The labelling was consider successful if the firing pattern of the neuron was modulated during current injection (i.e., to observe an increased activity during on-pulse and absence of activity in the off-pulse)^40,52^ and the process was stable for a minimum of 25 s followed by a rebound to spontaneous activity of the neuron after modulation. This procedure enabled us to map the recorded DMV neurons within the medulla. Thirty percent of recorded DMV neurons stopped firing either before or during the labelling procedure. In those cases the labelling protocol was carried on until the current reached 5-7 nA, in order to create a micro-trauma to verify the location within the DMV.

After recordings mice were euthanized (Nα-pentobarbital, 1.6 g/kg) and transcardially perfused with 4% paraformaldehyde, 15% picric acid in PBS, pH 7.4. The perfused brain was stored in PFA overnight and changed to 10% sucrose and 0.05% NaN3 solution for longterm storage.

### Slice physiology

Six weeks post vagal transfection, mice were deeply anesthetized with intraperitoneal injections of ketamine (200 mg/kg) – xylazine (23.32 mg/kg) and perfused transcardially with ice-cold modified artificial CSF (ACSF) oxygenated with 95% O_2_–5% CO_2_ and containing the following (in mM): 2.5 KCl, 26 NaHCO_3_, 1.25 Na_2_HPO_4_, 0.5 CaCl_2_, 10 MgSO_4_, 10 glucose, 0.4 Ascorbic acid, and 210 sucrose. The cerebellum, pons and medulla were rapidly removed, blocked in the coronal plane, and sectioned at a thickness of 240 μm in ice-cold modified ACSF. Slices were then submerged in ACSF, bubbled with 95% O_2_-5% CO_2_, and containing (in mM): 2.5 KCl, 126 NaCl, 26 NaHCO_3_, 1.25 Na_2_HPO_4_, 2 CaCl_2_, 2 MgSO_4_, and 10 glucose. The slices were transferred to the recording chamber mounted on an upright Zeiss Axioskop fixed-stage microscope and perfused with oxygenated ACSF at 32°C. A 60X, 0.9 NA water-immersion objective was used to examine the slice using standard infrared differential interference contrast video microscopy. Patch pipette resistance was typically 3-4.5 MΩ. A junction potential of 7-8 mV was not corrected. For cell-attached current-clamp recordings of firing patterns, whole-cell current clamp recordings and voltage clamp recordings of potassium currents the pipette contained (in mM): 135.5 KCH_3_SO_3_, 5 KCl, 2.5 NaCl, 5 Nα-phosphocreatine, 10 HEPES, 0.2 EGTA, 0.21 Na_2_GTP, and 2 Mg_1.5_ATP (pH=7.3 with KOH, 280-290 mOsm/kg). For whole cell voltage clamp recordings of calcium currents the pipette contained (in mM): 111 CsCH_3_SO_3_, 12.5 CsCl, 1 MgCl_2_, 0.1 CaCl_2_, 10 HEPES, 1 EGTA, 0.21 Na_2_GTP, and 2 Mg1.5ATP (pH=7.3 with CsOH, 280-290 mOsm/kg). In some experiments biocytin (0.2 % w/v) was added to this internal solution. We used a mixture of synaptic receptor blockers containing the following (in μm): 50 d-APV, 5 NBQX, 10 SR 95531, 1 CGP 55845, 10 mecamylamine, and 10 atropine. For whole cell calcium current recordings, HEPES-based bath solutions were used. The first solution contained (in mM): 137 NaCl, 1.8 CaCl_2_, 1 MgCl_2_, 5.4 tetraethylammonium (TEA)-Cl, 10 4-AP, 0.001 tetrodotoxin (TTX), 5 HEPES and 10 glucose (pH=7.3 with NaOH), and the second solution was identical except for an equimolar substitution of CaCl_2_ with CoCl_2_ (pH=7.3 with NaOH). To measure the effect of the Gi DREADD on the autonomous firing rate slice were incubated in 10 μM water-soluble CNO (HelloBio, UK). Kv4 channels were blocked with preincubation of the slices in 1.3 μM phrixotoxin-2, a highly selective Kv4 channel antagonist (Alomone, Jerusalem, Israel).

### Brain stem histology

The brain stem was sliced at 30 μm thickness with a cryostat (Leica CM1950) and the tissue was stored in an antifreeze solution^53^. For the immunofluorescent staining, after 4 x rinse/5 mins in 0.1M PB solution, the slices were incubated in normal donkey serum for 1 hour. Then, the tissue was incubated overnight in the primary antibodies dissolved with 0.3% Triton 0.1M PB with 10% normal donkey serum. The following primary antibodies were used: Mouse anti-human synuclein (1:5000, Thermo Fisher Scientific; RRID:AB_1954821; Recombinant rabbit anti-phospho S129 alpha synuclein, 1:2000, Abcam 51253);the goat anti-ChAT (1:100, Millipore; RRID: AB_262156)]. On the second day, the tissue was incubated in the secondary antibodies at the room temperature for 2 hours after washing steps. The following secondary antibodies were used: anti-mouse Alexa 488 (1:1000); anti-goat cy5 (1:1000); streptavidin conjugated to Cy3 (Abcam, UK); biotinylated donkey anti-rabbit (1:1000, Jackson ImmunoResearch). The sections treated with biotinylated antibodies were further incubated for one hour in avidin-biotin-peroxidase solution (ABC Elite, Vector laboratories, Burlingame, USA). The protein was finally visualized by a 0.1M PB solution containing 5% 3,3’-diaminobenzidine (DAB) (Sigma, Germany) and 0.02% H_2_O_2_. The sections were mounted on glass slides and dehydrated in a series concentration of ethanol followed an incubation of cresyl-violet solution (Sigma, Germany). Afterwards, the DAB/Nissl stained sections were shortly incubated in xylene (Merck, Germany) and coverslipped using mounting gel (Toluene solution, Fisher Scientific). The tissue of fluorescent staining was mounted (Vectashield, Vector Laboratories) on glass slides after 4x rinse in 0.1M PB.

The brain stem samples for the cohort of the juxtacellular recording were cut in to 50 μm thick slices (VT1000S microtome, Leica). Sections were rinsed in PBS and then incubated in a blocking solution (0.2 m PBS with 10% horse serum, 0.5% Triton X-100, 0.2% BSA). The following primary antibodies were used (in carrier solution, 22°C, overnight); monoclonal mouse anti-human α-synuclein (1:1000, 4B12, Thermo Fisher Scientific); the goat anti-ChAT (1:100, Millipore; RRID:AB_262156). The following secondary antibodies (in carrier solution, 22°C, overnight): AlexaFluor-488 donkey anti-goat, AlexaFluor-488 donkey antimouse and Streptavidin AlexaFluor-568 (1:1000; Molecular Probes) were used. Sections were rinsed in PBS and mounted on slides (Vectashield, Vector Laboratories).

For semiquantitative optical density analysis of the human α-synuclein expression in DMV neurons, we used ImageJ software (http://rsbweb.nih.gov/ij/) to determine the mean immunosignal intensities of ChAT-positive regions of interest in the brain stem slices. We used three confocal slices through the DMV from each mouse with a total of three AAV-A53T and AAV-EV injected mice for the analysis. Each confocal slice was a collapsed *z*-stack of 10 adjacent 1-μm-thick optical slices chosen so that they arose primarily from within individual DMV neurons. Thus, while it is not possible to rule out the possibility that some of the fluorescence arises from other processes (e.g., glia), the majority of the signal is confined to ChAT-immunopositive neuronal somatodendritic domains in the DMV.

### Stomach and intestine histology

After 200 nl AAV1/2-A53T-SNCA (the same virus used for vagal injection) intramuscularly injected in the stomach, two weeks after surgery, the mouse was euthanized for immunofluorescent staining. The staining procedure was processed as previously described^54^.

### Fluorescence microscopy and cell size analysis

#### Cell surface area

Confocal images of biocytin-filled cells reacted with streptavidin conjugated to Cy3 were captured on a Nikon confocal 1AR microscope under a Plan Apochromat Lambda 60x oil objective. Images were over-sampled in the z-plane for maximum preservation of 3D structure (pixel size xy = 0.138 μm; voxel depth z = 0.125 μm), and were processed and analyzed with FIJI software (ImageJ, NIH, version 2.0.0). The area encompassing the cell soma was cropped to discard neurites. A Gaussian blur was applied to the voxels (sigma radius = 1.0), and where necessary cells were filled so that only voxels on the cell surface were used for calculations. A threshold was set for each cell, and the total surface area were analyzed using the 3D objects counter plugin (FIJI). Only complete cells were included in the analysis.

#### Cell Diameter

6 weeks after injection with either AAV-EV (N=2 mice) or AAV-A53T (N=3 mice), we took confocal images of ChAT / human α-synuclein staining from 30 μm thick slides of DMV. In each slide, we collapsed 10-15 adjacent 1-μm-thick confocal images from the middle part of each slide in to a single image. We collected 5 such collapsed images that were approximately 150 μm apart from between AP: −7.32 to AP:−7.64 (from bregma). In each of the 5 images, we randomly selected ChAT-positive neurons with or without immunoactivity of anti-human α-synuclein from the medial-dorsal portion of DMV and measured the longest axis of their soma using the ImageJ software (NIH, USA).

### Stereological quantification of DMV neuron number

The number of Nissl-stained DMV neurons was stereologically quantified, as previously described^55^. Briefly, the DMV was delineated from every 5^th^ section of the DMV between Bregma −7.32 mm and −7.20 mm. Neuronal counts were performed on a Zeiss Axio Imager 2 microscope or an Olympus BX53, fitted with a Prior ProScan III motorised stage. The optical fractionator technique was used, and was facilitated by either Stereo Investigator version 10, (MBF Biosciences), or newCAST software (Visiopharm). Total cell numbers were calculated according to the methods outlined by Gundersen and Jensen^56^. A guard zone of at least 1 μm was set. Coefficient of error was calculated with values <0.10 accepted^56^.

### Data and Statistical Analysis

Data were analyzed and curve fitting was done using custom-made code on Matlab (The Mathworks, Natick, MA) software. The two-tailed Wilcoxon rank-sum test (RST) was used to test for changes of medians in two independent sample comparisons. The two-tailed Wilcoxon signed-rank test (SRT) was used to test for changes of medians in matched-paired comparisons. Two-tailed ANCOVA was used to test the difference between curves. The null hypotheses were rejected if the P values were below 0.05. Throughout the main manuscript and the supplementary data uppercase N’s (that are mentioned in the figures and figure captions) indicate the number of mice; and lowercase n’s (that appear in the figures) indicate the number of individual neurons recorded.

