## Supplementary Information for "α-Synuclein-induced Kv4 channelopathy in mouse vagal motoneurons causes non-motor parkinsonian symptoms"

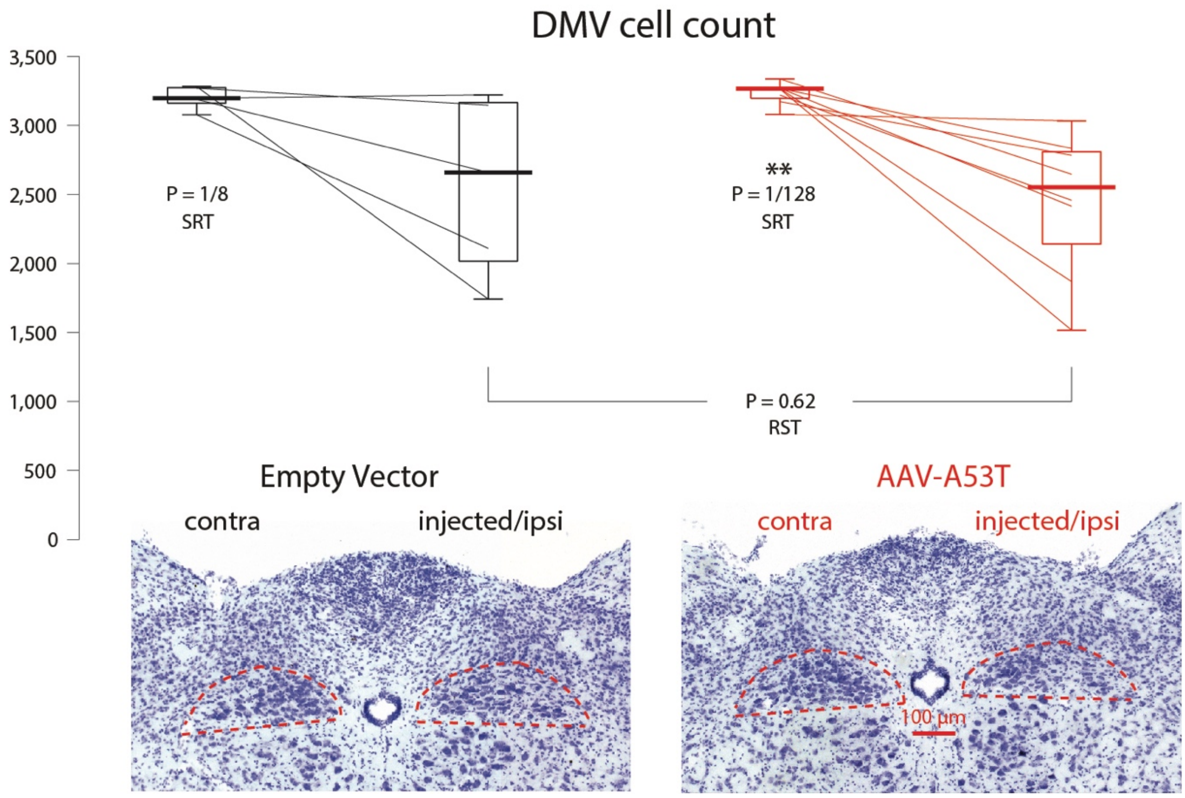


**Supplementary Figure 1. Variable DMV cell loss following cervical vagal AAV injections is not attributable to the α-synucleinopathy *per se*.** Box plot of the total unbiased stereological count of Nissl-stained neurons in the DMV in the noninjected (contralateral) and injected (ipsilateral) sides of mice injected with either AAV-EV (black) or AAV-A53T (red). Insets: examples of Nissl-stained slices of the dorsal medulla, with the DMV indicated by the dashed red line. RST – two-tailed Wilcoxon Rank-Sum test; SRT – Wilcoxon Signed-Rank test.

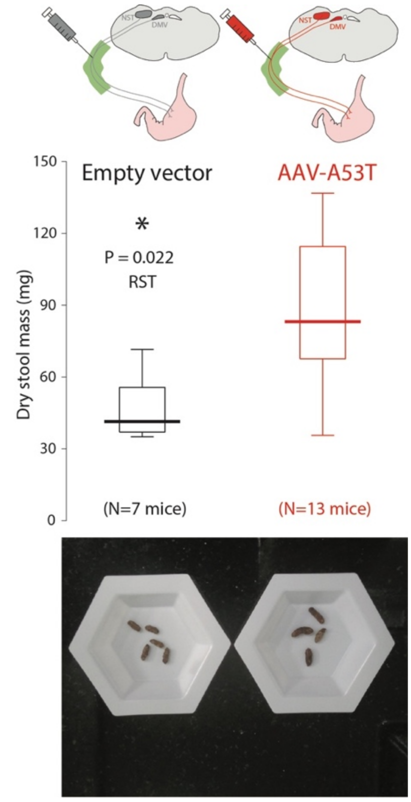


**Supplementary Figure 2. Adult onset medullary α-synucleinopathy increases mouse stool size.** Comparison of total dry stool mass between mice transfected with AAV-EV (left) or AAV-A53T (right). RST – two-tailed Wilcoxon Rank-Sum test.

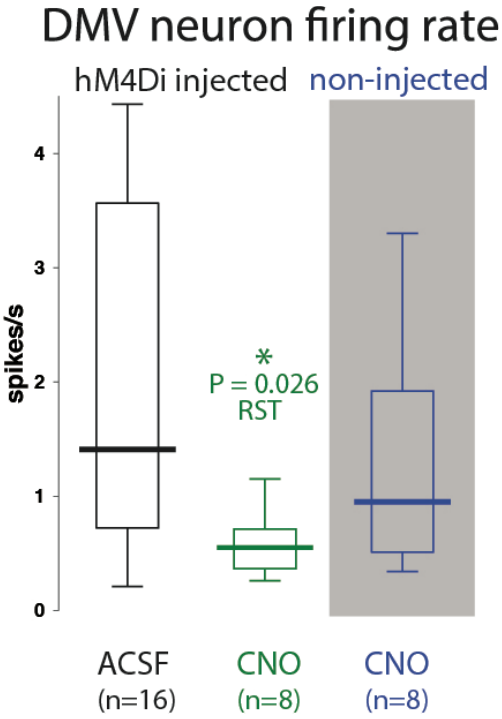


**Supplementary Figure 3. Chemogenetic inhibition of DMV neuron firing rate.** Recording of the spontaneous firing rate of DMV neurons in acute brain slices from ChaT-cre mice that underwent unilateral injection of AAVs harboring the hM4Di DREADD demonstrated that preincubation and superfusion of the slices in 10 µM CNO (N=2 mice) but not ACSF (N=3 mice) lowered the firing rate. Incubation of slices from non-injected hemispheres (N=2 mice) in CNO had no effect on the firing rate. RST – two-tailed Wilcoxon Rank-Sum test.

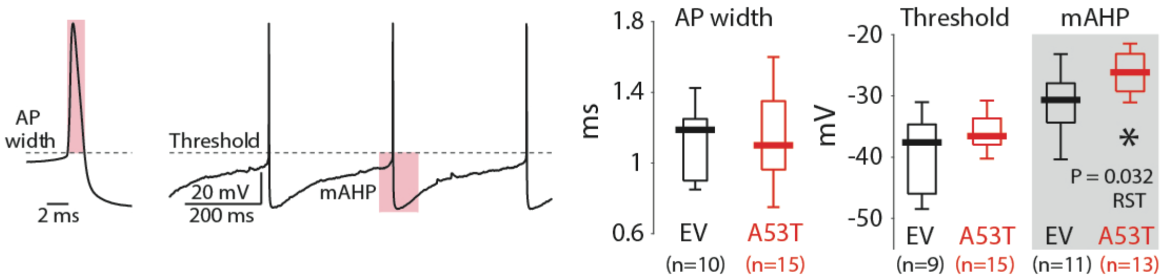


**Supplementary Figure 4. Action potential properties in DMV neurons are largely unchanged in the adult onset medullary α-synucleinopathy model.** Action potential (AP) width (width of pink box), threshold and afterhyperpolarization (AHP) amplitude (length of pink box, recorded as negative value relative to AP threshold) were measured and compared between mice transfected with AAV-EV (N=7 mice) and AAV-A53T (N=5 mice) (box plots). AP threshold was unchanged, ruling out changes in the persistent $Na^+$ current. AP width was unchanged ruling out changes in the large- conductance $Ca^{2+}$-activated $K^+$ (BK) current. AHPs were slightly reduced suggesting a possible reduction in the small conductance $Ca^{2+}$-activated $K^+$ (SK) current[1,2]. RST – two-tailed Wilcoxon Rank-Sum test.
